# A novel framework for modeling the evolution of cross-scale ecological assembly

**DOI:** 10.1101/2020.03.17.994319

**Authors:** Amanda S. Gallinat, William D. Pearse

## Abstract

Community assembly can be driven by species’ responses to environmental gradients, and interactions within (*e.g.*, competition) and across (*e.g.*, herbivory) clades. These ecological dynamics are mediated by species’ traits, which are in turn shaped by past evolution. As such, identifying the drivers of species assembly is made difficult by the differing temporal and spatial scales of ecological and evolutionary dynamics. Two recent advances have emerged to address the cross-scale challenge of modeling species assembly: phylogenetic generalized linear mixed modeling (PGLMM) and earth observation networks (EONs). PGLMM integrates through time by modeling the evolution of trait-based community assembly, while EONs synthesize across space by placing standardized site-level species occurrence data within their regional context. Here we describe a framework for combining these tools to investigate the drivers of species assembly, and so address three outstanding questions: (1) Does evolution adapt or constrain regional-scale environmental responses? (2) Do evolved responses to past competition minimize or enhance present-day competition? (3) Are species’ cross-clade associations evolutionarily constrained? We provide a conceptual overview of how PGLMM and EONs can be synthesized to answer these questions, and provide exemplar Bayesian PGLMM code. Finally, we describe the capacity of these tools to aid in conservation and natural resource management, including predicting future colonization by rare and invasive species, vulnerable mutualisms, and pest and pathogen outbreaks.

## 1 Bridging Ecology & Evolution: A Challenge of Scale

Evolution is the context for present-day ecological interactions, which in turn shape the evolution of species (Webb et al., 2002), although evolutionary biology and community ecology have proven notoriously difficult to integrate empirically. This has predominantly been an issue of scale; classically, ecological processes operate across smaller spatio-temporal scales than evolutionary processes (Webb et al. (2002); Cavender-Bares et al. (2009); but see, *e.g.*, Richardson et al. (2014)), but for practical reasons biodiversity data often represent *either* local *or* regional scales. Models and data sets that integrate across multiple spatial and temporal scales to capture processes dependent on dynamics that are both broad- and fine-scale in nature are difficult to generate and so rare. Yet, they are necessary to describe the interaction of evolutionary history and ongoing ecology in driving species occurrences over space and time (Pearse et al., 2018).

The lack of such cross-scale statistical models and empirical data has limited our ability to make broad inferences about the processes that determine the distribution of biodiversity. For instance, it is generally accepted that environmental filtering constrains species’ distributions at the regional scale, while at local scales competition and facilitation within and among clades (*e.g.*, vascular plants, grasses, mammals, or rodents) additionally constrain community assembly (Kraft et al., 2015). Yet it is also generally accepted that this is an over-simplification, and that disentangling the interaction of these niche-based processes requires the integration of regional and local data (Cadotte & Tucker, 2017). Even the same process can have profoundly different effects when operating at different scales. For instance, dispersal limitation can mediate coexistence and priority effects when operating at a finer spatio-temporal scale, while at broader scales it can drive speciation (Vellend, 2010).

The next great challenge for those who seek to understand and predict community structure is to solve this problem of parsing cross-scale processes by explicitly modeling those processes in cross-scale data (McGill, 2019). Recent efforts to address this challenge have highlighted the need for tools which incorporate biogeographic history and present-day ecological constraints to test evolutionary theory, interpret ecological patterns, and predict future community assemblages (Pearse et al., 2018). Here we describe how advances in ecological monitoring across spatial scales, and models incorporating phylogeny with ecological data, can be combined to model the evolution of ecological community assembly. Critically, we focus solely on methods and datasets that already exist, with the aim of catalyzing the generation of both empirical academic research and actionable insights for conservation and natural resource management.

## 2 New Tools for the Study of Community Assembly

While the last few decades have seen great conceptual advances in our understanding of what drives biodiversity (Chesson, 2000; Ricklefs, 2008; Vellend, 2010), the empirical application of these frameworks has proven difficult (with exceptions, of course, some of which we outline below). Here we describe how advances in data—Earth observation networks (EONs)—and statistics—a family of eco-phylogenetic models known as phylogenetic generalized linear mixed models (PGLMM; Ives & Helmus, 2011)—advance our understanding of the cross-scale evolutionary and ecological drivers of species assembly.

### 2.1 Earth Observation Networks

Empirically parsing how regional environmental and local within- and cross-clade processes motivate ecological assembly requires data sets that capture all three processes. Specifically, this challenge requires overlapping local-scale assemblage data for multiple clades that connect at the regional scale (see Table 1). Earth observation networks (EONs) like the USA’s National Ecological Observatory Network (NEON; Schimel et al., 2007) and Australia’s Terrestrial Ecosystem Research Network SuperSites (TERN; Karan et al., 2016) employ sampling designs that meet all of these requirements. The prescribed, top-down approach of these EONs pairs intentional site distribution with robust sampling protocols to capture regional environmental responses for a wide range of taxa. Nested within this regional site design, EONs generate local assemblage data which can be used to investigate the interplay between species historic and current niches through the phylogenetic patterns of co-occurrence within clades. Furthermore, EONs provide local-scale species assemblage data for *multiple* clades; the power to model the phylogenetic structure of co-occurrences across clades (*e.g.*, flowering plants and their insect pollinators, or ticks and their rodent hosts) is a singular contribution of EONs toward our understanding of the biotic drivers of species distributions. Only using data from EONs can these local-scale within- and cross-clade dynamics be modeled together, within the context of regional environmental filtering, to compare their relative contributions to species assembly.

**Table 1:** Data requirements for building complexity from *β*-trait models, to *α*-trait models, and finally to models of cross-clade associations. Focusing on three questions central to understanding the process of species assembly, we provide descriptions of the types of data required to address each question (including sampling breadth and density), examples of existing data sets, and a conceptual illustration of data collection. In the illustrations of data collection, each color represents a different species or taxon, depicted on the phylogenies at the right. Double arrows indicate closely-related species that have either evolved to minimize co-occurrence (top panel) or have constrained evolution such that they compete in the present (middle panel). At the regional scale (top panel), species within a clade (co-)occur based on environmental sensitivities that drive broad-scale distributions; the double arrow indicates species that have evolved not to co-occur. At the local scale (middle panel), species that overlap in distribution at the regional scale may not co-occur due to within-clade interactions; the double arrows indicate evolutionary constraints such that species compete in the present. Investigating cross-clade associations (bottom panel) requires that the regional- and local-scale data illustrated here above be collected for multiple clades; for species pairs across clades, the presence of one, both, or neither can be used to identify the phylogenetic patterns of associations.

| Question | Data Requirements | Examples | Data Collection |
| --- | --- | --- | --- |
| **(1)** Does evolution adapt or constrain regional-scale environmental responses? | Spatial extent includes an environmental gradient across which (co-)occurrence patterns vary<br>Assemblage sampling is sufficient to estimate phylogenetic structure of β-traits | Global Biodiversity Information Facility (GBIF: www.gbif.org; Edwards, 2000)<br>eBird (www.ebird.org; Sullivan et al., 2014)<br>Specimen data, e.g. US Integrated Digitized Biocollections (iDigBio: www.idigbio.org; Matsunaga et al., 2013) |  |
| **(2)** Do evolved responses to past competition minimize or enhance present-day competition? | Requirements for **(1)**<br>Sampling density within each local site captures variation in species co-occurrence patterns<br>Assemblage sampling is sufficient to estimate phylogenetic structure of α-traits | International Long-Term Ecological Research Network (ILTER: Mirtl et al., 2018)<br>U.S. Forest Inventory & Analysis (FIA: www.fia.fs.fed.us; Smith, 2002)<br>North American Breeding Bird Survey (BBS: www.pwrc.usgs.gov/bbs; Sauer et al., 2013) |  |
| **(3)** Are species' cross-clade associations evolutionarily constrained? | Requirements for **(1)** and **(2)**<br>Data overlap spatially and temporally for associated clades; interaction networks further inform associations of interest<br>Assemblage sampling is sufficient to estimate phylogenetic structure across (monophyletic) clades | U.S. National Ecological Observatory Network (NEON: www.neonscience.org; Schimel et al., 2007)<br>Australia's Terrestrial Ecosystem Research Network (TERN) SuperSites (supersites.tern.org.au; Karan et al., 2016) |  |

EONs have been criticized for lacking a central research question (*e.g.*, Lindenmayer et al., 2018), but implicit in any EON is a set of core hypotheses about scaling that support the synthesis of fields like macro-ecology, evolutionary biology, and community ecology (and are ideal for eco-phylogenetic analyses like PGLMM; see below). Despite being in their infancy, EONs have already informed our understanding of emergent properties of biodiversity (Guerin et al., 2019), trait variation (Bloomfield et al., 2018), and niche structure (Read et al., 2018). Read et al. (2018) are particularly notable for using EON data to re-examine, and extend, classic macro-ecological work about niche packing that had been conducted with bottom-up networks (*e.g.*, Ernest, 2005). Bottom-up networks synthesize existing sites’ data, and examples such as the US Long-Term Ecological Research network (LTER) have facilitated fundamental contributions to our understanding of biodiversity. For instance, Greenville et al. (2018) used LTER data to show that the impacts of climate change and wildfires have varied across biomes over the last 60 years, leveraging historic data that EONs, which are still in their infancy, would struggle to provide. Yet the homogenized data collection, management, and dissemination of EONs (Kao et al., 2012) permits more meaningful assessments of relative abundances and absences and allows for the kind of unprecedented cross-site synthesis that is essential to parsing the drivers of species assembly across scales. While we will focus on the advantages of EON data alone in our proposed framework, we predict that data from top-down EONs, bottom-up networks, as well as other types of networks (e.g. citizen science networks; see Table 1) will increasingly be combined, each benefiting from the complementary advantages of the other.

### 2.2 Phylogenetic Generalized Linear Mixed Models

Leveraging the unique sampling design of EONs to model cross-scale processes driving species assembly hinges on the availability of statistical tools that span evolutionary and ecological timescales. Eco-phylogenetic approaches incorporate the evolutionary history of species (their phylogeny) into analyses of current species distributions to determine how the shared biogeographic history of related species shapes their present-day ecology. The release of global phylogenies (*e.g.*, Faurby & Svenning, 2015) and automated tools (*e.g.*, Smith & Walker, 2018) have made the phylogenetic data required for such analyses available to ecologists.

Traditionally, eco-phylogenetic analyses have focused on whether co-occurring species are more or less closely-related than would be expected by chance. Under the assumption of niche conservatism (wherein closely-related species tend to resemble one-another), the co-occurrence of close relatives is the product of environmental filtering on shared tolerances, and distantly-related species co-occur through competitive exclusion on the basis of similar traits (Webb et al., 2002; Cavender-Bares et al., 2009). This simple, yet powerful, framework has driven advances in our understanding of how ecological structure varies across clades (Swenson et al., 2006), and how it interacts with the evolution of species’ traits (Rabosky et al., 2011; Pearse et al., 2019). Yet it has also led to the understanding that mapping eco-phylogenetic pattern onto particular ecological or evolutionary processes is fraught with difficulty (Mayfield & Levine, 2010); mapping pattern onto process is essentially impossible if we cannot contrast the relative strengths of differing drivers to explain observed patterns. Thus, recent advances in the field of eco-phylogenetics have aimed to synthesize deep- and shallow-time ecological and evolutionary processes into a single model to contrast the relative strength of potential drivers of biodiversity.

Phylogenetic generalized linear mixed models (PGLMM; Ives & Helmus, 2011) have emerged as a powerful way to quantify species’ occurrences or abundances as a function of species’ traits and environmental conditions (Box 1). PGLMM goes beyond describing the structure of assemblages (*e.g.*, containing closely-related, functionally distinct species) to describe the processes structuring assemblages (*e.g.*, divergent trait evolution followed by competition on that divergent trait). By using random effect terms to account for site- and species-level variation and measuring phylogenetic patterns among those random effects, PGLMM can be used to measure phylogenetically-patterned differences in how species interact with each other and their environment. Multiple power analyses of PGLMM have been published (Pearse et al., 2014; Ives & Helmus, 2011; Rafferty & Ives, 2013), therefore we focus here on new applications of this tested tool.

#### Box 1

Past and potential uses of PGLMM

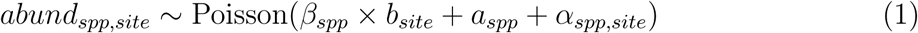

Equation 1 defines a PGLMM (Ives & Helmus, 2011), where *abund*_*spp,site*_ is the abundance of species (*spp*) across sites (*site*), *a*_*spp*_ is the overall abundances of species, and other terms are defined below. This is a PGLMM because phylogeny can inform its random effects, *a*_*spp,site*_ and *β*_*spp*_.

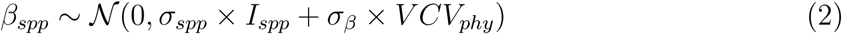

Equation 2 models the *β*-trait in two parts. The first accounts for each species’ independent variation—an identity matrix with dimensions equal to the number of species (*I*_*spp*_) multiplied by a scalar (*σ*_*spp*_)—the second accounts shared evolutionary history—the phylogenetic co-variance matrix (*V CV*_*phy*_) multiplied by a scalar (*σ*_*β*_). Thus 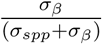 is Pagel’s *λ* (see Hadfield, 2010), testing for evolutionary constraint of species’ *β*-trait responses to the environment (*b*_*env*_). Alternative transformations of *V CV*_*phy*_ can be used to test for different evolutionary models.

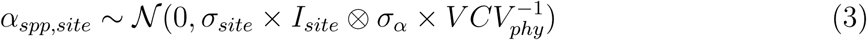

Equation 3 accounts for the *α*-trait in two parts. The first accounts for each species’ independent variation in site-level abundances—an identity matrix with dimensions equal to the number of sites (*I*_*site*_) multiplied by a scalar (*σ*_*site*_)—the second accounts for evolutionary repulsion of species’ co-occurrences—the inverse of the phylogenetic covariance matrix 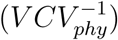 multiplied by a scalar (*σ*_*α*_). These terms are combined through their Kronecker product, and the magnitude of *σ*^*β*^ reflects the importance of repulsive trait evolution.

In the supplementary materials, we show how to fit these models to empirical data, and also how subtle changes to these equations can test the hypotheses we outline in the main text in sections 3.1, 3.2, and 3.3. These include extending this model to account for the phylogenetic structure of another clade (the basis of section 3.3).

The PGLMM framework is well-suited to analyzing cross-scale patterns within EON data because it integrates broad-scale processes underlying the macro-evolution of species traits with their local-scale consequences for community assembly. Indeed, PGLMM often *requires* a combination of regional and local community data, since the composition of nearby sites rarely encompasses sufficient environmental variation for macro-scale conclusions. Studies of small assemblages, or large assemblages within small regions, are insufficient to detect both environmental filtering and the evolutionary signal of environmental filtering (Swenson et al., 2006).

## 3 Three cross-scale questions PGLMM and EONs can answer

Together, EONs and PGLMM provide the scale of data and statistical flexibility necessary to investigate three cross-scale processes driving species distributions: the evolution of (1) abiotic constraints, (2) within-clade interactions, and (3) cross-clade associations. To address these research themes, we propose a synthetic framework that integrates differing mechanisms of the macro-evolution of species’ traits with both macro- and local-scale assembly processes based upon those traits (Figure 1). In this framework, species traits evolve either *adaptively* (constrained by broad optima; but see Pearse et al. (2018)) or *repulsively* (reducing similarity among close-relatives, putatively to minimize competition; see Nuismer & Harmon, 2015). These same traits are then filtered by the regional environment and drive local-scale competition to dictate assemblage composition. Associations such as pollination and trophic dynamics constitute the cross-clade environment which can further constrain species occurrences on the basis of resource availability, mutualism, and pressure from predators, parasites, and pathogens. Reflecting an increasing recognition of the role of species’ associations in community assembly, this framework incorporates cross-clade co-occurrence within the context of within-clade and regional dynamics to understand how species’ evolution and community assembly shape, and are shaped by, these processes. Below we describe the utility of this framework for leveraging EON data to test the contributions and mechanisms of abiotic, within-clade, and cross-clade processes. We also describe how this approach can be extended to provide support for the conservation and management of vulnerable species, interactions, and communities.

**Figure 1:**
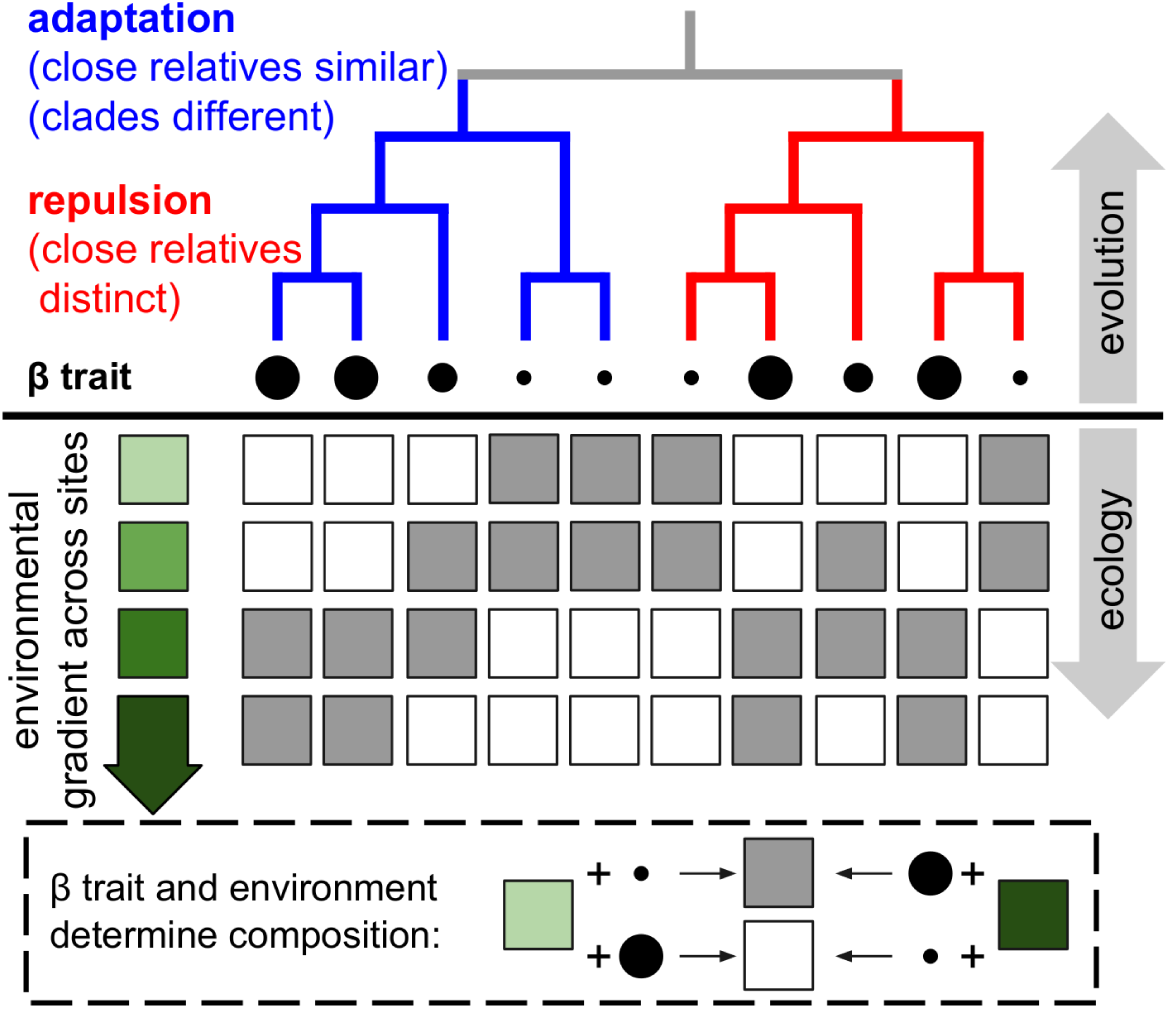
Conceptual overview of regional-scale and macro-evolutionary hypotheses that can be tested by analyzing EON data with PGLMM. Ten species (columns) and their presence in four sites along an environmental gradient (rows) are shown. The match of species’ *β* traits (circles) to the environment (green boxes)—environmental filtering—determines community membership. The *β*-traits themselves are shown as they would be if evolving under two distinct macro-evolutionary models: repulsion (the red clade on the right) and adaptation (the blue clade on the left).

A large part of our framework is based around the concepts of *β*- and *α*-traits, which parse the differing roles that species traits can play in community assembly (Silvertown et al., 2006). Regional habitat affiliations are determined by *β*-traits, which regulate species’ tolerances to environmental filters and are hypothesized to evolve early and be strongly conserved through time. By contrast, *α*-traits evolve later, and determine local-scale co-occurrence, competition, and environmental filtering. Drought tolerance, for instance, is considered a *β*-trait at the regional scale, with soil moisture explaining differences among habitat types and thus regionally clustering related species with shared water use traits (Moeslund et al., 2013). Yet such tolerance is an *α*-trait when driving local-scale hydrological niche differentiation in a different system (Silvertown et al., 2015). That the same trait can be subject to different evolutionary and ecological forces in different clades and environments is unsurprising. Indeed, we argue below that modeling traits in this fashion will allow us better generalize seemingly discordant results across systems and unpick the cross-scale drivers of community assembly.

Our framework is not hypothetical, and can be applied by combining readily-available tools. To demonstrate this, in the supplementary materials we provide examples of how to fit PGLMM to address each of the conceptual processes outlined below. This code maps directly onto the overview given in Box 1.

### 3.1 Does evolution adapt or constrain regional-scale environmental responses?

While it is now well-established in the comparative literature that assessing patterns in species’ traits without accounting for their shared evolutionary history can introduce bias (Freckleton et al., 2002), it is only beginning to be recognized that the same biases can affect studies of species’ ecological assembly. For instance, PGLMM has been shown to reveal even stronger signals of trait-based assembly by accounting for phylogenetic non-independence (Li & Ives, 2017). As in the comparative literature, however, phylogeny can play a much greater role in our understanding of community assembly than as a mere statistical correction. At the regional scale, phylogeny can be used to investigate the evolutionary constraints on, and evolutionary processes underlying, species’ environmental responses. Many mechanisms are invoked to explain the pattern of phylogenetic signal in *β*-traits, many of which involve the environment. It follows that a goal of species assembly modeling should be to measure the strength of niche conservatism itself as reflected in species’ environmental responses, in which the responses themselves are *β*-traits. By incorporating random effects for species and sites, PGLMM can be used to model species’ environmental responses 1) as trait-based environmental tolerances, using the interaction of species’ measured traits and the environment, 2) as unmeasured (latent) traits for environmental responses, estimated using phylogeny (as described in Box 1), and 3) as coefficients describing species-specific responses to environmental gradients. In the latter case, environmental responses are still modeled as latent traits, but can now be used to estimate the extent to which those latent traits are better when informed by phylogeny, and to test for evolutionary constraints on environmental sensitivity. Critically, within this approach latent traits are *β*-traits, and measure environmental responses. By virtue of phylogeny, this method can also be used to estimate the evolutionary signal of sensitivities and predict those for species lacking data (Swenson, 2013).

Beyond patterns of environmental response among species, a deeper understanding of the mode by which trait-environment relationships develop is critical to understanding how *β*-traits affect species assembly. If functional traits are driving environmental responses, PGLMM can be used to co-estimate the evolution and ecology of those traits, allowing for more nuanced inference of each component. Techniques such as divergence-order time analysis can also be used to determine which came first: evolutionary constraint on the trait, or environmental niche. Equally, application of PGLMM allows us to take advantage of the recent explosion of macro-evolutionary hypotheses and methods, and it is now common to contrast Brownian motion (a null model of conserved evolution; see Losos, 2011) with Ornstein-Uhlenbeck (OU—a model of adaptive evolution; see Butler & King, 2004) and accelerating/decelerating models of trait evolution (Blomberg et al., 2003) using AIC-based model selection (Boettiger et al., 2012). OU models may describe the adaptive landscape of macro-evolution (Harmon & Uyeda, 2014), but it remains to be seen whether their macro-evolutionary trait trade-off surfaces apply in the literal landscapes of EON data. As described in Box 1, differing models could be fit to EON data through linear transformations of species’ phylogenetic variance-covariance matrices (see Freckleton et al., 2002) to contrast different rates and modes of evolution for niches and traits.

Such combined eco-phylogenetic techniques offer a solution to a pervasive challenge in the study of trait macro-evolution: the mapping of phylogenetic pattern onto process. It is well-known that phylogenetic signal, and/or support for a particular statistical model of trait evolution, does not always reveal the mechanism of evolution. For example, statistical support for Brownian motion could be the result of neutral drift or environmental constraint (Revell et al., 2008). Here again, however, the breadth of data from EONs combined with the statistical flexibility of PGLMM provide a way forward. By modeling the evolution of species’ environmental responses (*β*-traits), we can explicitly link ecological mechanism to evolutionary model, and thus use statistical means to address mechanistic questions. For example, the hypothesis that a trait’s pattern of phylogenetic signal results from environmental constraint can be tested with a PGLMM that quantifies both evolutionary constraint and environmental filtering in the present. We have a limited amount of past data from which to infer evolutionary process, but in principle can collect infinite data in the present to inform our understanding of the past.

### 3.2 Do evolved responses to past competition minimize or enhance present-day competition?

There is still great debate over how we can best measure species’ niche differences, and so estimate the magnitude of competition (D’Andrea & Ostling, 2016). While some clades have candidate *α*-traits known to impact local-scale niche structure (*e.g.* specific leaf area in vascular plants, Cornwell & Ackerly (2009)), the latent trait approach described in section 3.1 can also aid in the discovery of traits relevant for competition. PGLMM can go further by measuring not just how species’ *α*-traits drive local-scale assembly, but also jointly estimating how those traits have evolved. Specifically, PGLMM can be used to model species occurrence as a function of trait dispersion, phylogenetic dispersion, both, or—most critically for examining modes of *α*-trait evolution—trait dispersion combined with some evolutionary model of the trait. The latter need not be limited to simple competitive exclusion models of competition based around niche differences (see Mayfield & Levine, 2010); PGLMM could be applied to more complex models of fitness and stabilizing niche differences (see, for example, Chesson, 2000). Future efforts could also incorporate other explicit models of competition (*e.g.* Drury et al. (2016) and Morlon et al. (2016)) in much the same way we suggest using phylogenetic transformations to test different models of (latent) trait evolution in section 3.1. Just as phylogenetic conservatism can mask patterns of *β*-trait assembly, so too could it for *α*-traits, but the combination of EON data and PGLMM offers a unique opportunity to model complex local-scale species assembly processes within the regional context of environmental filtering, potentially on the same traits.

The traditional picture of excluding competition driving local-scale ecological assembly has been further complicated by an increasing understanding of the role of Neutral processes (Hubbell, 2001; Vellend, 2010). Within Neural theory, there is a growing disconnect between two classes of model. In the first class, ecological assembly within clades is random with respect to species’ traits, and so stabilizing forces are absent and species do not compete for resources (Hubbell, 2001). In the second, traits drive ecological assembly but the *evolution* of those traits is neutral; stabilizing forces and competition are present and act on randomly evolved traits (Rosindell et al., 2015). To examine the validity of the first class, sufficient environmental variation is required to detect differences across species’ ranges, as provided by EONs, while the second class requires the integration of ecological and evolutionary dynamics provided by PGLMM.

Rather than evolving to compete on the basis of similar traits in the present, species may also have evolved to reduce ecological competition. While within macro-ecology such evolution, as manifest as character displacement, is reasonably uncontroversial (Brown & Wilson, 1956; Schluter & McPhail, 1992), in community ecology it is controversial to invoke evolved responses to past competition, or the ‘ghost of competition past’, to explain an absence of present-day competition (Connell, 1980). The integration of evolutionary modeling into community ecological models through PGLMM offers the opportunity to not only directly test for this ecological ghost, but also improve macro-evolutionary models of species traits. While new and promising methods are being developed to measure and simulate the repulsive evolution of species’ traits (*e.g.*, Drury et al., 2016), it is difficult to distinguish among different models on the basis of purely extant species’ traits (Nuismer & Harmon, 2015). As with the combined modeling of *β*-traits, we suggest the solution is to model the evolution of species’ interaction coefficients within PGLMM. Only such joint eco-evolutionary models can allow us to distinguish between whether traits have evolved to minimize competition, or whether evolutionary constraint has forced species to compete in the present.

### 3.3 Are species’ cross-clade associations evolutionarily constrained?

The *α*- and *β*-trait based model of ecological assembly described above, like many filtering-based models of community assembly, ignores interactions across clades. However, species’ distributions result not just from their own intrinsic cross-scale evolutionary and ecological dynamics, but also from their interactions with other groups. In the name of tractability, community ecologists often taxonomically restrict their assemblages (Ricklefs, 2008; Vellend, 2010) and interaction ecologists are frequently forced to overlook temporal and spatial variation in species’ interactions (Poisot et al., 2015). The co-diversification of groups like plants and pollinators or herbivores and mammals and their parasites have been rich and controversial fields of study. Even the most specialized mutualisms that form classic models of diversification have provided surprisingly little evidence for pure, strict coevolution by cospeciation (Hembry & Althoff, 2016). And while the timing of diversification in many clades has been well-studied, there are fewer studies of the evolution of the traits driving co-occurrences and associations among these clades (but see, for example, Rafferty & Ives, 2013). Thus, modeling the role of the cross-clade environment in shaping communities and driving biodiversity is likely to provide answers to many outstanding evolutionary questions.

For species to interact, they must first be capable of co-occurring in the same space, and thus a natural starting place for modelling variation in network structure is the *α*/*β* framework outlined above. Using this framework, the simplest model of co-occurrence across clades is a statistical interaction between each group’s parameters. While species’ responses to environmental drivers are often phylogenetically patterned, the pattern of responses to other species is unclear (Cavender-Bares et al., 2009). The PGLMM framework provides a solution by estimating the effect of the cross-clade environment on species occurrences. By comparing data describing the presence of two interacting species, as in figure 2, co-occurrence data can be modeled as they would be in a PGLMM of interaction network data (Rafferty & Ives, 2013), incorporating two separate phylogenies to generate expected pairwise interaction rates. Clades’ responses and effects could both be phylogenetically patterned (top left of figure 2), neither could be phylogenetically patterned (bottom right of figure 2), or only the effect *or* response could be patterned (top right and bottom left of figure 2), resulting in observable differences in overlapping distribution. These differences can provide insight into the contrasting drivers of species assembly for interacting clades; for instance, the phylogenetic patterns of co-occurring hosts may be a strong driver of parasite community assembly, while host assembly may be predominantly driven by environmental limits and within-clade dynamics. This method therefore builds on intra-clade regional environmental and local-scale species assembly models to enable researchers to estimate the drivers and future of cross-clade interactions.

**Figure 2:**
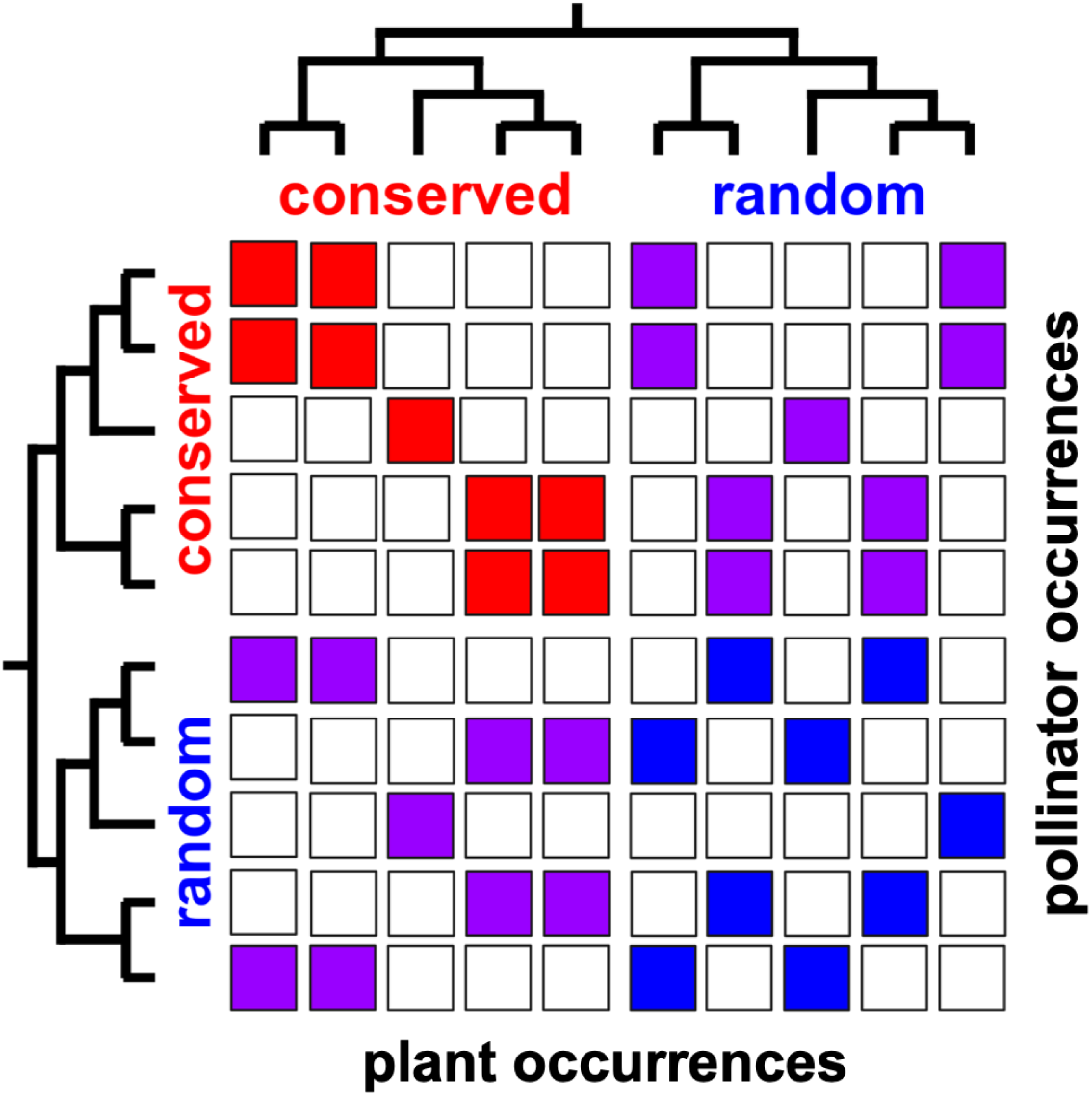
Cross-clade hypotheses of phylogenetic co-occurrence, shown here as plants and pollinators, that can be tested with PGLMM. Ten plant (columns) and ten pollinator (rows) species are shown, with recorded co-occurrences between them shown with colored cells. For each group two clades, each with either phylogenetically conserved or random co-occurrences, are shown. Each clade combination represents a different hypothesis about plant–pollinator co-occurrences.

Central to the concept of the cross-clade environment is the idea that species evolve to match the environment of other species, and vice versa. Species’ traits drive many interactions, including those among predators and prey, parasites and their hosts, and mutualists (Werner & Peacor, 2003). The PGLMM method improves our ability to test the effect of traits on species co-occurrence (as for species assembly) through comparison among models describing separate (for each species) and combined regional and local trait- and environment-based drivers, employing the same model transformations described in sections above to test for adaptive, repulsive, and neutral evolutionary process. For instance, this framework can be used to test if species’ interactions thought to drive plant structure are evolutionarily conserved or divergent (Cavender-Bares et al., 2009), or whether species’ interactions are too diffuse to permit co-evolution (Zillio et al., 2005). Inherent to this analysis is the need for sufficient spatial breadth of species assemblage data to capture both regional and local drivers, as well as a broad sample of where species do and do not co-occur.

## 4 Applying the PGLMM framework to current and future conservation challenges

Phylogeny has already proven an informative tool for conservation planning (*e.g.*, Isaac et al., 2007), restoration (Hipp et al., 2015), and trait prediction (Swenson et al., 2017). Combining EON data with PGLMM adds an additional dimension with which to formulate niche-based conservation approaches as related to environmental response, competitive interactions, and cross-clade associations. The framework we outline can be used to rank species in terms of their likelihood of successful colonization and/or interaction with other members of a community. Below we outline some of the ways in which such a prioritization could be produced and used.

### Environmental responses (*β*-traits)

PGLMM allows for the prediction of whether rare, under-studied, or invasive species could colonize a region on the basis of phylogeny, with applications for restoration and invasive species management. For instance, if species’ environmental responses are phylogenetically patterned, or if closely related species tend to (or tend not to) co-occur, a site’s environmental characteristics and current species assemblage can be used to assess how successful an introduced or invasive species will be there. Using these tools, the impact of invasive species could be predicted even before species have arrived on a continent.

### Competitive interactions (*α*-traits)

Darwin’s theory of naturalization predicts that invasive species are less likely to establish near native congeneric species due to higher competition and vulnerability to local pests and pathogens (Daehler, 2001). Thus, in observable communities invasives species are expected to be both phylogenetically and functionally distant from native neighbors; however this expectation has proven somewhat controversial in practice (*e.g.*, Thuiller et al., 2010). The PGLMM framework can be used to test the theory of naturalization across a broad range of communities and taxa. Furthermore, using freely available functional trait and phylogenetic (or at least taxonomic) data, and the estimated parameters from the PGLMM described above, it is possible to rank the likelihood of species successfully invading sites to help managers plan for invasions.

### Cross-clade associations

Joint evolutionary and ecological modeling of interaction networks could aid in the prediction of pest and pathogen outbreaks, vulnerable mutualisms, and shifting trophic dynamics. PGLMM can be used to determine whether species have co-diversified or whether similar environmental constraint has forced them to associate. In the former case, we might expect limited host-switching and spill-over under climate change, whereas in the latter case we might expect more. Previous research identifying phylogenetic patterns in spillover among hosts (Parker et al., 2015) indicates the potential for eco-phylogenetic tools to combine the phylogenetic likelihood of host spillover with the ecological limitations of pests and pathogens as well as their hosts to predict future associations. Understanding when species are evolutionarily constrained not to co-occur will allow efforts to be targeted at those species who are only held back by ecological opportunity, which, in a changing world, may not be the case for long.

## 5 Conclusion

Central to the framework described here is the integration of evolutionary and ecological tools, theory, and their differing strengths to identify cross-scale drivers of species assembly. In macro-evolutionary biology, the patterns of species’ traits fit statistical models well, but determining underlying mechanisms is hard. We simply cannot run evolutionary experiments over millions of years. A strength of ecology, on the other hand, is the ability to examine mechanisms at the local scale, but that scale limits the potential to address underlying generative processes (Peters et al., 2014). The PGLMM approach bridges these gaps, strengthening evolutionary techniques by providing more data (EONs) to test mechanisms, and broadening ecological scope with the generality required to predict across regions. Opportunities to combine PGLMM and EONs to integrate across evolution and ecology will almost certainly continue to expand. Increasingly sophisticated macro-evolutionary models are only going to need more data with which to test their predictions, and as ecology becomes more predictive the temporal and spatial scale over which models will need to operate can only grow. PGLMM provides a way forward to merge both fields for mutual benefit, but require datasets that would have been implausible before EONs made them available to all.

## Supporting information

Supplemental PGLMM R Code

## Acknowledgments

The authors wish to thank Matthew Helmus for comments on an earlier version of the manuscript. This work was funded by National Science Foundation grant EF1802605.

## Biosketch

Amanda S. Gallinat’s research bridges ecology and evolution to understand how environmental change affects biodiversity, phenology, and ecosystem function. William D. Pearse focuses on the use of phylogenies to infer how ecological assembly and function operate and on the role of phylogenies in conservation prioritization.

## Data Accessibility Statement

No new data are released as part of this manuscript. R code for Bayesian PGLMM is included in the Supporting Information.

