## Supplemental PGLMM R Code for "A novel framework for modeling the evolution of cross-scale ecological assembly": Gallinat_Pearse_PGLMM-Supplement.R

```
########################################
########################################
# Supplement to: A novel framework for 
#   modeling the evolution of
#   cross-scale ecological assembly

# Amanda S. Gallinat and William D. Pearse
########################################
########################################

# Some general notes:

# * Ignore 'divergent transitions' warnings. These are nothing to
#   worry about in these models, but, as with all Bayesian models, you
#   should perform model diagnostic checks (e.g., posterior predictive
#   checks) before using the output of a model. See Gelman et
#   al. (2013; Bayesian Data Analysis) for more details.
# * We use 'set.seed' to make the simulations below completely
#   reproducible. Since this makes all random numbers in your R
#   session the same, you may wish to restart R if doing further
#   simulation/analysis work.
# * To make the Bayesian posteriors reproducible, we also specify a
#   'seed' in each stan call. This is not something that would require
#   you to re-start your computer inbetween runs, but is probably
#   something that you wouldn't want to use in your own analyses.
# * You may still get slightly different (but qualitatively the same)
#   results if using an old verison of R (<3.5; released in July
#   2019). Later versions of R randomly permute data slightly
#   differently to address a known bug ("significant user-visible
#   changes in 3.6.0";
#   https://cran.r-project.org/doc/manuals/r-devel/NEWS.html).
# * To make these models fit in a reasonable timeframe on a laptop, we
#   work with fairly simply datasets (in particular for Question
#   2.2). By all means use this code as a starting point for more
#   complex simulations and analyses.
# * Running "options(mc.cores=XXX)", where XXX is the number of
#   threads your computer can run (2 or 4 on most laptops) will make
#   the model fitting *much* faster
# * The code below will install packages if they are missing from your
#   computer, which means it will take longer to run the first time.

########################################
########################################
# R packages and functions #############
########################################
########################################
# Load / install packages
if(!require("geiger")){
    install.packages("geiger", dependencies=TRUE); library(geiger)}
if(!require("mvtnorm")){
    install.packages("mvtnorm", dependencies=TRUE); library(mvtnorm)}
if(!require("pez")){
    install.packages("pez", dependencies=TRUE); library(pez)}
if(!require("lme4")){
    install.packages("lme4", dependencies=TRUE); library(lme4)}
if(!require("rstan")){
    install.packages("rstan", dependencies=TRUE); library(rstan)}

########################################
# Community simulation functions #######
########################################
# Inverse logit function
inv.logit <- function(x)
    exp(x) / (exp(x)+1)
# Environmental filtering
make.env.comm <- function(trait, env, occupancy, tree, rescale=TRUE){
    data <- expand.grid(species=names(trait), site=names(env), stringsAsFactors=FALSE)
    data$trait <- trait[data$species]
    data$env <- env[data$site]
    data$occupancy <- occupancy[data$species]
    data$num_species <- as.numeric(gsub("s","",data$species))
    data$num_sites <- as.numeric(factor(data$site))
    data$prob <- data$trait * data$env + data$occupancy
    if(rescale)
        data$prob <- inv.logit(data$prob)
    data$presence <- rbinom(nrow(data), 1, data$prob)
    return(data)
}
# Scramble competition
make.comp.comm <- function(comp.trait, n.spp, env, tree, scale=2){
    # Make overdispersed VCV from trait
    vcv <- as.matrix(dist(comp.trait))
    vcv <- max(vcv) - vcv
    vcv <- diag(length(env)) %x% solve(vcv)
    
    # Make overdispersed VCV from phylogeny
    # vcv <- solve(vcv(tree))/scale
    # vcv <- diag(length(env)) %x% vcv

    # Draw presences/absences
    pres.abs <- matrix(inv.logit(rmvnorm(1, sigma=vcv)), nrow=length(env), byrow=TRUE)
    for(i in seq_len(nrow(pres.abs)))
        pres.abs[i,] <- ifelse(pres.abs[i,]>=sort(pres.abs[i,], decreasing=TRUE)[sample(n.spp,1)], 1, 0)
    comm <- matrix(as.numeric(t(pres.abs)), nrow=length(env), ncol=length(comp.trait), dimnames=list(names(env), tree$tip.label), byrow=TRUE)
        
    # Format data and return
    data <- comparative.comm(tree, comm, traits=data.frame(trait=comp.trait), env=data.frame(env=env))
    data <- as.data.frame(data)
    data$site <- factor(data$site)
    data$species <- factor(data$species)
    data$num_species <- as.numeric(gsub("s","",data$species))
    data$num_sites <- as.numeric(factor(data$site))
    return(data)
}
# Association simulation
make.assoc <- function(base.data, base.tree, base.trait, partner.trait){
    # Make prob. of association matrix
    assoc <- 1 - plnorm(abs(outer(base.trait, partner.trait, FUN=`-`)))
    
    # Add in partner interactions on basis of probabilities
    partner.comm <- matrix(0, nrow=nrow(base.data), ncol=length(partner.trait))
    for(i in seq_len(nrow(base.data))){
        if(base.data$presence[i]==1)
            partner.comm[i,] <- as.numeric(runif(length(partner.trait)) <= assoc[base.data$num_species[i],])
    }

    # Reformat data and return
    data <- base.data[rep(seq_len(nrow(base.data)), length(partner.trait)),]
    names(data)[c(6,8,9)] <- c("num_base_species","base_prob","base_presence")
    data$num_partner_species <- rep(seq_along(partner.trait), each=nrow(partner.comm))
    data$partner_presence <- as.numeric(partner.comm)
    return(data)
}

########################################
########################################
# Simulate data used throughout ########
########################################
########################################
# Basics
set.seed(1234)
n.spp <- 50
tree <- sim.bdtree(n=n.spp, seed=1234)
n.sites <- 50
site.names <- apply(as.matrix(expand.grid(letters,letters,stringsAsFactors=FALSE)), 1, paste, collapse="")[seq_len(n.sites)]
env.sites <- setNames(rnorm(n.sites), site.names)

# Overall OCCupancies (intercepts) and traits (slopes) under three
#   different models of evolution: CONStrained (phy.signal), LABile
#   (phylogenetically random), and COMpetition (repulsion)
cons.occ <- sim.char(tree, .05, root=0)[,,1]
lab.occ <- setNames(sample(cons.occ), names(cons.occ))
comp.occ <- setNames(rmvnorm(1, rep(0,n.spp), solve(vcv(tree))/10)[1,], tree$tip.label)
cons.trait <- sim.char(tree, .05, root=0)[,,1]
lab.trait <- setNames(sample(cons.trait), names(cons.trait))
comp.trait <- setNames(rmvnorm(1, rep(0,n.spp), solve(vcv(tree))/10)[1,], tree$tip.label)

# Make datasets based around traits to exemplify biologically plausible scenarios
cons.env <- make.env.comm(cons.trait, env.sites, lab.occ, tree)
# - Constrained evolution of species' environmental responses,
#   environmental assembly on the basis of those traits, and
#   phylogenetically neutral overall occupancies
ghost.comp <- make.env.comm(comp.trait, env.sites, lab.occ, tree)
# - Evolution to minimise competition (the 'ghost of competition
#   past') with environmental assembly on the basis of those traits,
#   and phylogenetically neutral overall occupancies
s.tree <- drop.tip(tree, 16:50)
cons.comp <- make.comp.comm(cons.occ[1:15], c(2,5), env.sites[1:15], s.tree)
# - Constrained evolution of species' environmental responses, and
#   competitive (scramble) assembly on the basis of those same traits
# NOTE: we're now working with fewer sites, because these competition
#   models are quite computationally intensive and we would like these
#   examples to run in a reasonable timeframe


# Make association dataset
base.n.spp <- 10; partner.n.spp <- 10
base.tree <- sim.bdtree(n=base.n.spp, seed=2345); partner.tree <- sim.bdtree(n=partner.n.spp, seed=3456)
assoc.n.sites <- 10; assoc.env.sites <- setNames(rnorm(assoc.n.sites), letters[seq_len(assoc.n.sites)])
base.env.trait <- sim.char(base.tree, .05, root=0)[,,1]; base.occ <- setNames(sample(sim.char(base.tree, .05, root=0)[,,1]), names(base.env.trait))
base.assoc.trait <- sim.char(base.tree, .05, root=0)[,,1]; partner.assoc.trait <- sim.char(partner.tree, .05, root=0)[,,1]
# - First, make a "dummy" dataset with a 'base' species and a
#   'partner' associated species, but have them be independent. This
#   is to provide a test of what happens when two species are independent
base.env <- make.env.comm(base.env.trait, assoc.env.sites, base.occ, base.tree)
# - Now simulate a "base" species and a "partner" species, where the
#   partner is dependent on the base species (i.e., is found in
#   association with that base species)
assoc.data <- make.assoc(base.env, base.tree, base.assoc.trait, partner.assoc.trait)

########################################
########################################
# Q1: Environmental responses  #########
########################################
########################################

########################################
# Model 1.1: presence ~ trait * env ####
one.one <- glmer(presence ~ env * trait + (1|species), family=binomial, data=cons.env)
summary(one.one)
# - Significant interaction between environment and trait. Note the random
#   effect accounts for species-level pseudoreplication; if desired
#   you can add in site-level terms too but for this toy example we do
#   not think they are necessary

########################################
# Model 1.2: presence ~ trait * env ####
#                 WITHOUT phylogeny ####
# - First, let's fit it using frequentist methods
one.two <- glmer(presence ~ (env|species), family=binomial, data=cons.env)
#NOTE: Sometimes, you may get a boundary fit warning/error. If so, try
#      the fit again (perhaps with a different random number seed or
#      optimiser). This is a difficult model to numerically fit.
summary(one.two)

# Extract the latent trait (the coefficients of the interaction term),
# then re-order to match the simulated trait and plot
latent.traits <- setNames(ranef(one.two)$species$env, rownames(ranef(one.two)$species))
latent.traits <- latent.traits[names(cons.trait)]
plot(latent.traits ~ cons.trait)
abline(0,1, col="red", lwd=2)
cor.test(latent.traits, cons.trait)
# ... Not a perfect reconstruction (remember, there is error added in
#     the simulation) but not bad either

# - Second, let's do this using Bayesian methods. Go through the
#   comments below *slowly* because everything else builds on this
one.two.stan.code <- "
// Tell rstan what our data are
data{
    int Ntotal;              // Number of rows in our dataset
    int Nspp;                // Number of species
    int presence[Ntotal];    // The response variable (presence/absence)
    real env[Ntotal];        // An explanatory variable (environment)
    int spp[Ntotal];         // An explanatory variable (species)
}
// The parameters/coefficients we want to estimate and report back
parameters{
    vector[Nspp] spp_intercepts;  // Each species' overall occupancy
    vector[Nspp] spp_slopes;      // Each species' environmental response
}
// Some calculations rstan is going to perform internally to help model-fitting
transformed parameters{
    vector[Ntotal] predictions;   // Make a variable to hold our model predictions
    // Loop over all our input data and specify our model, which is:
    // a species' intercept (overall occupancy) +  env response x the environment
    for (i in 1:Ntotal)
        predictions[i] = spp_intercepts[spp[i]] + spp_slopes[spp[i]]*env[i];
}
// Fit our model to our predictions
model{
    // Species' occupancies and responses are drawn from uninformative priors
    spp_intercepts ~ normal(0, 1); 
    spp_slopes ~ normal(0, 1);
    // The model itself: our presences are drawn from our predictions
    presence ~ bernoulli_logit(predictions);
}
"
one.two.stan.model <- stan(model_code=one.two.stan.code,
                data=list(Ntotal=nrow(cons.env), Nspp=n.spp,
                presence=cons.env$presence,
                env=cons.env$env, spp=cons.env$num_species),
                iter=1000, chains=4, seed=123456)

# Now let's match the output up (in much the same way as before, only
#   now using 'extract' to get the information from the posterior
#   distribution of our rstan model fit)
latent.traits <- apply(extract(one.two.stan.model)$spp_slopes, 2, median)
names(latent.traits) <- paste0("s", seq_along(latent.traits))
latent.traits <- latent.traits[names(cons.trait)]
plot(latent.traits, cons.trait)
cor.test(latent.traits, cons.trait)
# ... and, again, we've reconstructed the trait

########################################
# Model 1.3: presence ~ trait * env ####
#                 WITH phylogeny    ####
# Below is a modified Bayesian version of the model, with the changes
#   from model 1.3 highlighted
one.three.stan.code <- "
data{
    int Ntotal;
    int Nspp;
    int presence[Ntotal];
    real env[Ntotal];
    int spp[Ntotal];
    //    
    matrix[Nspp,Nspp]Vphy;     // Give the phylogeny as data
}
parameters{
    vector[Nspp] spp_intercepts;
    vector[Nspp] spp_slopes;
    // Coefficients for the phylogenetically-derived variance in model terms
    real<lower=0,upper=100> lam_intercepts;    // (with priors specified too) 
    real<lower=0,upper=100> lam_slopes;        // 
    // Coefficients for the NON-phylogenetically-derived variance in model terms
    real<lower=0,upper=100> null_intercepts;       // (also with priors)
    real<lower=0,upper=100> null_slopes;           //
}
transformed parameters{
    vector[Ntotal] predictions;
    for (i in 1:Ntotal)
        predictions[i] = spp_intercepts[spp[i]] + spp_slopes[spp[i]]*env[i];
}
model{
    // Now we draw our species coefficients to measure the importance of phylogeny
    spp_intercepts ~ multi_normal(rep_vector(0,Nspp), diag_matrix(rep_vector(null_intercepts,Nspp)) + lam_intercepts*Vphy); 
    spp_slopes ~ multi_normal(rep_vector(0,Nspp), diag_matrix(rep_vector(null_slopes,Nspp)) + lam_slopes*Vphy);
    //
    presence ~ bernoulli_logit(predictions);
}
"
one.three.stan.model <- stan(model_code=one.three.stan.code,
                data=list(Ntotal=nrow(cons.env), Nspp=n.spp,
                presence=cons.env$presence,
                env=cons.env$env, spp=cons.env$num_species, Vphy=vcv(tree)),
                iter=2000, chains=4, seed=123456)

# First thing to note: we can now get estimates for the latent traits
# and occupancies (we could have done this before, but there was no
# need). We're going to use a wrapper function for this now, to make
# this code easier to read and generalise.
.grab.latent.trait <- function(model, name, known.trait){
    latent <- apply(extract(model)[[name]], 2, mean)
    names(latent) <- paste0("s", seq_along(latent))
    return(latent[names(known.trait)])
}
cor.test(.grab.latent.trait(one.three.stan.model, "spp_slopes", cons.trait), cons.trait)
cor.test(.grab.latent.trait(one.three.stan.model, "spp_intercepts", lab.occ), lab.occ)
# ... great, nice positive correlations (i.e., we accurately estimated the traits)

# Now let's check for phylogenetic signal!
lam.slopes <- mean(extract(one.three.stan.model)[["lam_slopes"]])
null.slopes <- mean(extract(one.three.stan.model)[["null_slopes"]])
lam.slopes / (null.slopes + lam.slopes)
# ...the estimated phylogenetic signal (Pagel's Lambda) of the
# estimated environmental response is ~.7 (i.e., there's signal)
# (see Hadfield & Nakagawa (2010) J. Evol. Biol. 23(3): 494-508 for
# this method of calculating it)

# ...and let's do the same for the occurrence, using a wrapper
# function (again, to make the code easier to read and generalise)
.pagel.lam <- function(lam, null, model){
    phy <- mean(extract(model)[[lam]])
    null <- mean(extract(model)[[null]])
    return(phy / (phy + null))
}
.pagel.lam("lam_intercepts", "null_intercepts", one.three.stan.model)
# ... while phylogenetic signal isn't absent, as we modeled
# it to be, it is certainly weaker here than in examples where
# we have simulated signal. And if we explore the distribution 
# of intercepts... 
hist(extract(one.three.stan.model)[["lam_intercepts"]])
# ... values are centered at zero (but note that they cannot
# be less than zero because they are estimates of variance)

# (An aside: the latent trait we estimated in Q1.2 does have
# phylogenetic signal too, albeit when estimated using a different
# method)
fitContinuous(tree, latent.traits, model="lambda")
# (Look at the 'lambda' parameter in the output above)

########################################
########################################
# Q2: Competition  #####################
########################################
########################################

########################################
# Model 2.1: the ghost of competition past
two.one.stan.code <- "
data{
    int Ntotal;
    int Nspp;
    int presence[Ntotal];
    real env[Ntotal];
    int spp[Ntotal];
    matrix[Nspp,Nspp] Vphy;
    matrix[Nspp,Nspp] inv_Vphy; // The inverse of the phylogeny
    //                             (used to measure repulsive evolution)
}
parameters{
    vector[Nspp] spp_intercepts;
    vector[Nspp] spp_slopes;
    real<lower=0,upper=100> lam_intercepts;
    real<lower=0,upper=100> lam_slopes;
    real<lower=0,upper=100> null_intercepts;
    real<lower=0,upper=100> null_slopes;
    // Coefficients for the inverse-phylogenetically-derived variance in model terms
    real<lower=0,upper=100> inv_intercepts;       // (with priors specified too) 
    real<lower=0,upper=100> inv_slopes;           // 
}
transformed parameters{
    vector[Ntotal] predictions;
    for (i in 1:Ntotal)
        predictions[i] = spp_intercepts[spp[i]] + spp_slopes[spp[i]]*env[i];
}
model{
    // Now we draw our species coefficients to measure the importance of phylogeny *and its inverse*
    spp_intercepts ~ multi_normal(rep_vector(0,Nspp), diag_matrix(rep_vector(null_intercepts,Nspp)) + lam_intercepts*Vphy + inv_intercepts*inv_Vphy); 
    spp_slopes ~ multi_normal(rep_vector(0,Nspp), diag_matrix(rep_vector(null_slopes,Nspp)) + lam_slopes*Vphy + inv_slopes*inv_Vphy); 
    presence ~ bernoulli_logit(predictions);
}
"
two.one.stan.model <- stan(model_code=two.one.stan.code,
                           data=list(Ntotal=nrow(ghost.comp),
                                     Nspp=max(ghost.comp$num_species),
                                     presence=ghost.comp$presence,
                                     env=ghost.comp$env, spp=ghost.comp$num_species,
                                     Vphy=vcv(tree), inv_Vphy=solve(vcv(tree))),
                           iter=1000, chains=4, seed=123456)

# Let's compare the relative support for species' traits evolving
# under Brownian motion (lam_slopes), under repulsion (inv_slopes -
# the inverse of the phylogenetic covariance matrix), or null (no
# phylogenetic signal at all) using a wrapper function not unlike the
# one we used above to get at Pagel's lambda
.rel.support <- function(model, ...){
  terms <- list(...)
  vars <- setNames(numeric(length(terms)), names(terms))
  for(i in seq_along(terms))
      vars[i] <- median(extract(model)[[terms[[i]]]])
  return(setNames(vars / sum(vars), unlist(terms)))
}
.rel.support(two.one.stan.model, "lam_slopes", "inv_slopes", "null_slopes")
# ... repulsion most likely (these numbers sum to one: ~68% support
#     for repulsion, ~28% for null, but definitely not Lambda (~2%))

.rel.support(two.one.stan.model, "lam_intercepts", "inv_intercepts", "null_intercepts")
# ... contrasting with the intercepts, which were labile (~80%
#     support), and show no evidence of repulsion.
# ... note that we are very much at the limit of what we can do with
#     these kinds of models (this is exploratory work!) - the inverse
#     of the phylogenetic covariance matrix has an unusual
#     distribution of entries, and this makes model-fitting (and
#     simulation) difficult


# Model 2.2: competition in the present
two.two.stan.code <- "
data{
    int Ntotal;
    int presence[Ntotal];
    matrix[Ntotal,Ntotal] InvPhySite;   // Matrix of phylogeny x sites to detect competition
}
parameters{
    real<lower=0.0001, upper=10> null_sites;       // Null variance of sites
    real<lower=0.0001, upper=10> inv_sites;        // Phylogenetic repulsion of sites
    vector[Ntotal] predictions;
}
model{
    predictions ~ multi_normal(rep_vector(0,Ntotal), diag_matrix(rep_vector(fabs(null_sites),Ntotal)) + fabs(inv_sites)*InvPhySite);
    presence ~ bernoulli_logit(predictions);
}
"

two.two.stan.model <- stan(model_code=two.two.stan.code,
                           data=list(Ntotal=nrow(cons.comp),
                                     Nspp=max(cons.comp$num_species),
                                     Nsite=max(cons.comp$num_site),
                                     presence=sample(cons.comp$presence),
                                     spp=cons.comp$num_species,
                                     InvPhySite=((solve(vcv(s.tree))/10) %x% diag(max(cons.comp$num_site)))),
                           iter=2000, chains=4, seed=123456)
# NOTE: We are working with a matrix with as many rows and columns as
#       we have data entries - this is a computationally difficult
#       model to fit (be patient and careful with its output)
# Let's see the support for phylogenetic repulsion in these data
inv.sites <- extract(two.two.stan.model)[["inv_sites"]]
sum(inv.sites > .2) / length(inv.sites)
#...~95% of the posterior density is above 0.2, which we would argue
#   is reasonably strong support for phylogenetic overdispersion. Note
#   that there are many alternative ways of testing for this (this
#   approach is similar to that in Ives & Helmus 2011 ), and we have
#   used an unhelpful (uniform) prior here.
# NOTE: We are using very little data here (only 15 species) - play
#       around with larger datasets to see better performance, and
#       also (if you like) change to simulating using phylogeny (and
#       not a two-step process of phylogeny --> trait --> assembly) to
#       increase statistical power.

########################################
########################################
# Q3: Species' associations  ###########
########################################
########################################

########################################
# Model 3.1: presence ~ env * phylogeny + other.species
# (Contrast this model 1.3)
three.one.stan.code <- "
data{
    int Ntotal;
    int Nspp;
    int presence[Ntotal];
    real env[Ntotal];
    int spp[Ntotal];
    matrix[Nspp,Nspp]Vphy;
    //
    int Npartner;                              // # of partner species (c.f., Nspp)
    int partner[Ntotal];                       // Give partner presence as data
    matrix[Nspp,Nspp]VpartnerPhy;              // Partner phylogeny
}
parameters{
    vector[Nspp] spp_intercepts;
    vector[Nspp] spp_slopes;
    vector[Nspp] partner_intercepts;                    // Allow for partner presence/absence to predict base species' presence/absence
    real<lower=0,upper=100> lam_intercepts;
    real<lower=0,upper=100> lam_slopes;
    real<lower=0,upper=100> null_intercepts;
    real<lower=0,upper=100> null_slopes;
// Coefficients for the partner-derived variance in model terms
    real<lower=0,upper=100> lam_partner_intercepts;     // (phylogenetic component)
    real<lower=0,upper=100> null_partner_intercepts;    // (non-phylogenetic)
}
transformed parameters{
    vector[Ntotal] predictions;
    for (i in 1:Ntotal)
        predictions[i] = spp_intercepts[spp[i]] + spp_slopes[spp[i]]*env[i] + partner_intercepts[partner[i]]*partner[i];
    // Notice extra term added at the end for the partner (analogous to spp_intercepts)
}
model{
    spp_intercepts ~ multi_normal(rep_vector(0,Nspp), diag_matrix(rep_vector(null_intercepts,Nspp)) + lam_intercepts*Vphy); 
    spp_slopes ~ multi_normal(rep_vector(0,Nspp), diag_matrix(rep_vector(null_slopes,Nspp)) + lam_slopes*Vphy);
    partner_intercepts ~ multi_normal(rep_vector(0,Npartner), diag_matrix(rep_vector(null_partner_intercepts,Npartner)) + lam_partner_intercepts*VpartnerPhy);
    // Notice the intercepts term is the same as the above, but now we address the partner species
    presence ~ bernoulli_logit(predictions);
}
"

# First fit: as a null, let's model the 'base' species that doesn't
# depend on the partner species, to see what finding no association
# looks like
three.one.stan.model.base <- stan(model_code=three.one.stan.code,
                data=list(Ntotal=nrow(assoc.data), Nspp=base.n.spp,
                presence=assoc.data$base_presence,
                env=assoc.data$env, spp=assoc.data$num_base_species, Vphy=vcv(base.tree),
                Npartner=partner.n.spp, partner=assoc.data$num_partner_species,
                VpartnerPhy=vcv(partner.tree)),                
                iter=1000, chains=4, seed=123456)

# Test the relative importance of the partner on the base species' presences
round(.rel.support(three.one.stan.model.base, "null_intercepts", "lam_intercepts", "null_partner_intercepts", "lam_partner_intercepts"), 2)
# ... very low values for the partner terms (essentially 0) - this
#     species' presences isn't determined by the other species

########################################
# Now let's repeat the model fitting, but for the *other* species
#   ('partner') that is driven by the presence of the 'base' species
three.one.stan.model.partner <- stan(model_code=three.one.stan.code,
                data=list(Ntotal=nrow(assoc.data), Nspp=partner.n.spp,
                presence=assoc.data$partner_presence,
                env=assoc.data$env, spp=assoc.data$num_partner_species, Vphy=vcv(partner.tree),
                Npartner=base.n.spp, partner=assoc.data$num_base_species,
                VpartnerPhy=vcv(base.tree)),                
                iter=1000, chains=4, seed=123456)
# -  notice that, above, we've swapped "partner" and "base" around,
#    otherwise the data input is the same

round(.rel.support(three.one.stan.model.partner, "null_intercepts", "lam_intercepts", "null_partner_intercepts", "lam_partner_intercepts"), 2)
# ... now we can see the impact of partner on species' distributions,
#     explaining ~95% of the variance (70% + 25%)

.pagel.lam("lam_partner_intercepts", "null_partner_intercepts", three.one.stan.model.partner)
# ... and we can see that it has some phylogenetic signal (as it
#     should, because we simulated it to) - it is remarkable we can
#     detect this given we have only 10 species (in each clade) in
#     this dataset!
```
